# TaxAss: Leveraging a Custom Freshwater Database Achieves Fine-Scale Taxonomic Resolution

**DOI:** 10.1101/214288

**Authors:** Robin R. Rohwer, Joshua J. Hamilton, Ryan J. Newton, Katherine D. McMahon

**Affiliations:** Environmental Chemistry and Technology Program, University of Wisconsin-Madison; Department of Bacteriology, University of Wisconsin-Madison; School of Freshwater Sciences, University of Wisconsin-Milwaukee; Department of Civil and Environmental Engineering, University of Wisconsin-Madison

## Abstract

Taxonomy assignment of freshwater microbial communities is limited by the minimally curated phylogenies used for large taxonomy databases. Here we introduce TaxAss, a taxonomy assignment workflow that classifies 16S rRNA gene amplicon data using two taxonomy reference databases: a large comprehensive database and a small ecosystem-specific database rigorously curated by scientists within a field. We applied TaxAss to five different freshwater datasets using the comprehensive Silva database and the freshwater-specific FreshTrain database. TaxAss increased the percent of the dataset classified compared to using only Silva, especially at fine-resolution family-species taxa levels, while across the freshwater test-datasets classifications increased by as much as 11-40 percent of total reads. A similar increase in classifications was not observed in a control mouse gut dataset, which was not expected to contain freshwater bacteria. TaxAss also maintained taxonomic richness compared to using only the FreshTrain across all taxa-levels from phylum to species. Without TaxAss, most organisms not represented in the FreshTrain were unclassified, but at fine taxa levels incorrect classifications became significant. We validated TaxAss using simulated amplicon data with known taxonomy and found that 96-99% of test sequences were correctly classified at fine resolution. TaxAss splits a dataset’s sequences into two groups based on their percent identity to reference sequences in the ecosystem-specific database. Sequences with high similarity to sequences in the ecosystem-specific database are classified using that database, and the others are classified using the comprehensive database. TaxAss is free and open source, and available at www.github.com/McMahonLab/TaxAss.

**IMPORTANCE:** Microbial communities drive ecosystem processes, but microbial community composition analyses using 16S rRNA gene amplicon datasets are limited by the lack of fine-resolution taxonomy classifications. Coarse taxonomic groupings at phylum, class, and order level lump ecologically distinct organisms together. To avoid this, many researchers define operational taxonomic units (OTUs) based on clustered sequences, sequence variants, or unique sequences. These fine-resolution groupings are more ecologically relevant, but OTU definitions are dataset-dependent and cannot be compared between datasets. Microbial ecologists studying freshwater have curated a small, ecosystem-specific taxonomy database to provide consistent and up-to-date terminology. We created TaxAss, a workflow that leverages this database to assign taxonomy. We found that TaxAss improves fine-resolution taxonomic classifications (family, genus and species). Fine taxonomic groupings are more ecologically relevant, so they provide an alternative to OTU-based analyses that is consistent and comparable between datasets.

## INTRODUCTION

Microbial communities form the foundations of all ecosystems, yet interpretation of community data is limited by the difficulty of comparing across datasets. With the rapid development of massively parallel sequencing technology, scientists are increasingly able to fingerprint microbial communities using amplicon sequencing of marker genes such as the 16S rRNA gene. The resulting sequences are typically grouped into Operational Taxonomic Units (OTUs) defined by sequence identity or sequence variants. Comparison between amplicon datasets is difficult because OTUs are specific to each analysis. For clarity, this paper refers to 16S rRNA gene amplicon sequencing datasets as “datasets” and defines OTUs as a dataset’s sequence unit of measure, irrespective of whether those units represent clustered sequences, sequence variants, or unique sequences.

### Taxonomy Allows Cross-Study Analyses

OTUs are widely used to represent ecologically coherent entities (1), however they represent study-specific phylotypes that cannot be compared between datasets. Many common OTU definitions including sequence identity-based clustering (2), minimum entropy decomposition (3), and distribution-based clustering (4) are specific to each analysis, resulting in arbitrary OTU names. OTU definitions based on exact sequences, such as DADA2’s denoising approach (5) or defining OTUs as unique sequences, are still specific to the amplicon region and sequencing platform used in each study. For these reasons, direct comparison of OTUs between multiple datasets is most often impossible.

Taxonomic naming systems allow comparisons between datasets by creating consistent terminology and consistent phylogeny-determined boundaries between organisms. However, taxonomic naming is most useful when sequences can be classified to a fine level (e.g. family, genus, or species). Many abundant taxa have poorly resolved fine-scale phylogenetic structures in reference taxonomy databases (hereafter “databases”), resulting in only coarse classifications for large proportions of amplicon datasets (e.g. phylum, class, or order). Coarse taxonomic groupings often include diverse organisms with differing ecological roles, so analyses at coarse taxa levels miss underlying ecological dynamics (6). Fine-resolution taxonomic names are required to bridge the gap between ecologically relevant OTU-based analyses and consistent, comparable taxonomy-based analyses.

### Ecosystem-Specific Taxonomy Databases

Microbial ecologists from diverse sub-fields have created fine-resolution reference taxonomies by curating databases specific to their ecosystems. These ecosystem-specific databases are small compared to the large comprehensive databases compiled by Greengenes (7), Silva (8), and the Ribosomal Database Project (9), but they are generally well-curated with more finely resolved phylogenies for ecosystem-specific lineages. Examples of ecosystems with curated databases include the human oral cavity (10), the cow rumen (11), the honey bee gut (12), the cockroach and termite gut (13), activated sludge (14), and freshwater lakes (15). Ecosystem-specific databases are created to establish consistent vocabulary for common uncultured bacteria, create monophyletic reference structures, incorporate new reference information, and understand what the “typical” organisms are in a given ecosystem. Additionally, ecosystem-specific databases can be used to assign taxonomy to a finer resolution than can be achieved with a large comprehensive database.

### The FreshTrain

This paper demonstrates TaxAss’s efficacy using a variety of freshwater amplicon datasets, the comprehensive Silva database (8), and the ecosystem-specific Freshwater Training Set (FreshTrain) (15). The FreshTrain database was created in 2012 and was originally curated alongside Greengenes. FreshTrain versions match Greengenes and Silva at the phylum, class, and order levels, but at finer taxonomic levels the FreshTrain is curated based on additional information such as the geographical distribution of sequences. These finer levels are referred to as lineage, clade, and tribe and approximate the Linnaean family, genus, and species (15). The FreshTrain is available online at www.github.com/McMahonLab/TaxAss.

### Taxonomy Assignment Algorithm

Classification algorithms assign taxonomic names to OTUs based on their similarity to reference sequences in a database. The most commonly used classification algorithm was developed by Wang *et. al* (16) for the Ribosomal Database Project and is implemented in both mothur (17) and QIIME (18). This naïve Bayesian classifier (hereafter “Wang classifier”) assigns taxonomy to OTUs based on 8-mer signatures and reports a bootstrap confidence estimate for each assignment (16). This bootstrap confidence value is based on the repeatability of the OTU’s assignment with subsampled 8-mers, not on an absolute similarity measure. In a large database an OTU dissimilar to any reference sequences will not be classified repeatably as any one taxon, resulting in a low bootstrap confidence. However, in a small database an OTU dissimilar to any reference sequences nevertheless can be classified repeatably because there are fewer references from which to choose. We refer to this pitfall as “misclassification” when OTUs are classified as unrelated organisms and “overclassification” when OTUs are classified to a finer taxa level than warranted.

### Introducing TaxAss

We aimed to obtain fine-level taxonomy classifications in freshwater datasets by leveraging the ecosystem-specific FreshTrain database, while at the same time maintaining the full biological diversity of each dataset. To this end, we developed an open source taxonomy assignment workflow (TaxAss) that uses the popular Wang classifier as implemented in mothur and employs both an ecosystem-specific database and a comprehensive database. TaxAss maintains taxonomic richness and accuracy by only classifying OTUs that share high percent identity with ecosystem-specific reference sequences using the ecosystem-specific database. The remaining OTUs are classified using the comprehensive database. TaxAss scripts and step-by-step directions are available online at www.github.com/McMahonLab/TaxAss.

## RESULTS

### Methods Summary

TaxAss uses both an ecosystem-specific database and a large comprehensive database to improve taxonomic assignment resolution while maintaining richness. To classify the maximum possible number of OTUs and avoid forcing inaccurate ecosystem-specific classifications onto OTUs, the amplicon dataset is split into two groups using blastn prior to classification: OTUs with high percent identity to ecosystem-specific reference sequences and OTUs with low percent identity to ecosystem-specific reference sequences. The two groups are then classified separately using the Wang classifier and the appropriate database (Figure 1).

**Figure 1.**
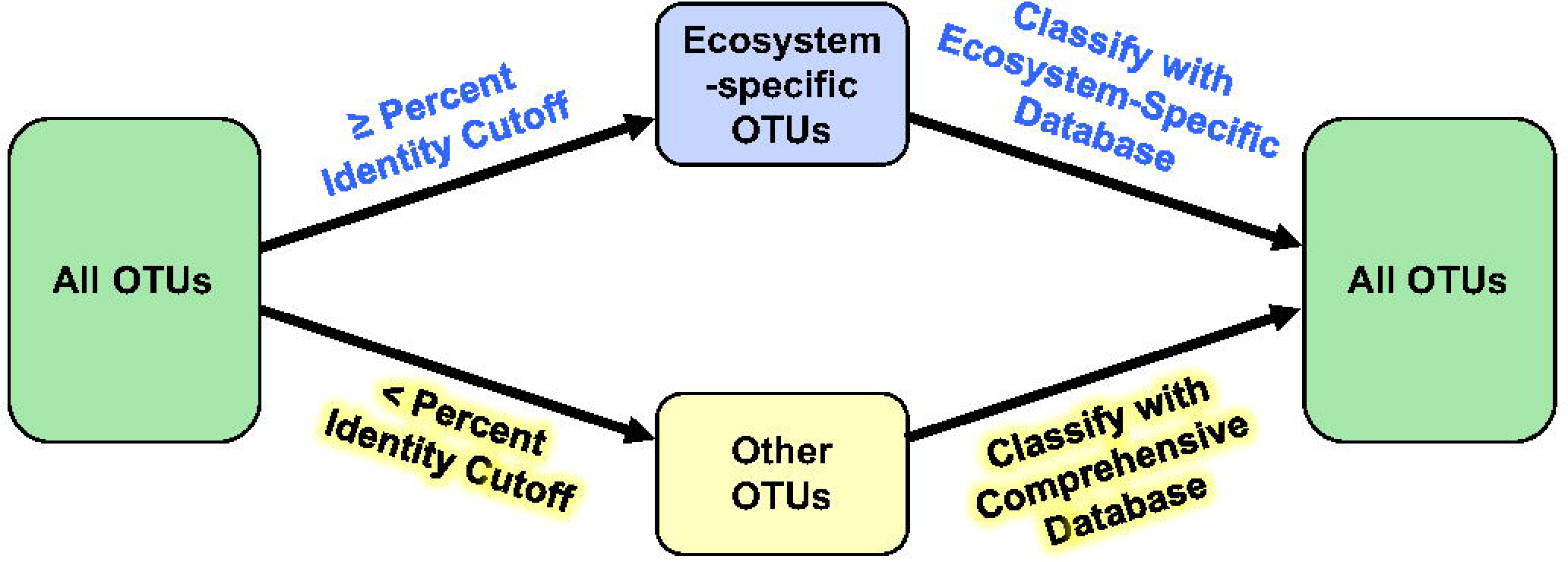
TaxAss Conceptual Diagram. TaxAss separates OTUs into two groups that are classified separately and then recombined. OTUs similar to any ecosystem-specific reference sequences are classified using the ecosystem-specific database, otherwise they are classified by the comprehensive database. BLAST is used to split the OTUs into groups (left arrows), and the Wang classifier is used to assign taxonomy (right arrows).

To test TaxAss we used SILVA version 132 and the FreshTrain as the comprehensive and freshwater-specific databases. We classified six 16S rRNA gene amplicon (tag) datasets spanning five freshwater ecosystems and a non-freshwater control. Sequences in these tag datasets are not directly comparable because they cover five different amplicon regions. We defined OTUs as the unique sequences remaining after basic quality filtering and chimera checking.

### Assignment Accuracy

To test the accuracy of TaxAss’s taxonomy assignments, we compared TaxAss results to a ground truth determined by manual alignment of full-length 16S rRNA gene sequences. For this test we used a full-length freshwater clone library dataset from Marathonas Reservoir, Greece (19), that was not previously incorporated into the FreshTrain. We manually aligned these full-length sequences to the FreshTrain, and then simulated a tag dataset by trimming the full-length sequences to the commonly used primer regions V4, V4-V5, and V3-V4. We classified this simulated tag dataset using TaxAss with the FreshTrain and Silva and compared the results to the ground truth results provided by manual full-length alignments and phylogenetic analysis (Figure 2).

**Figure 2.**
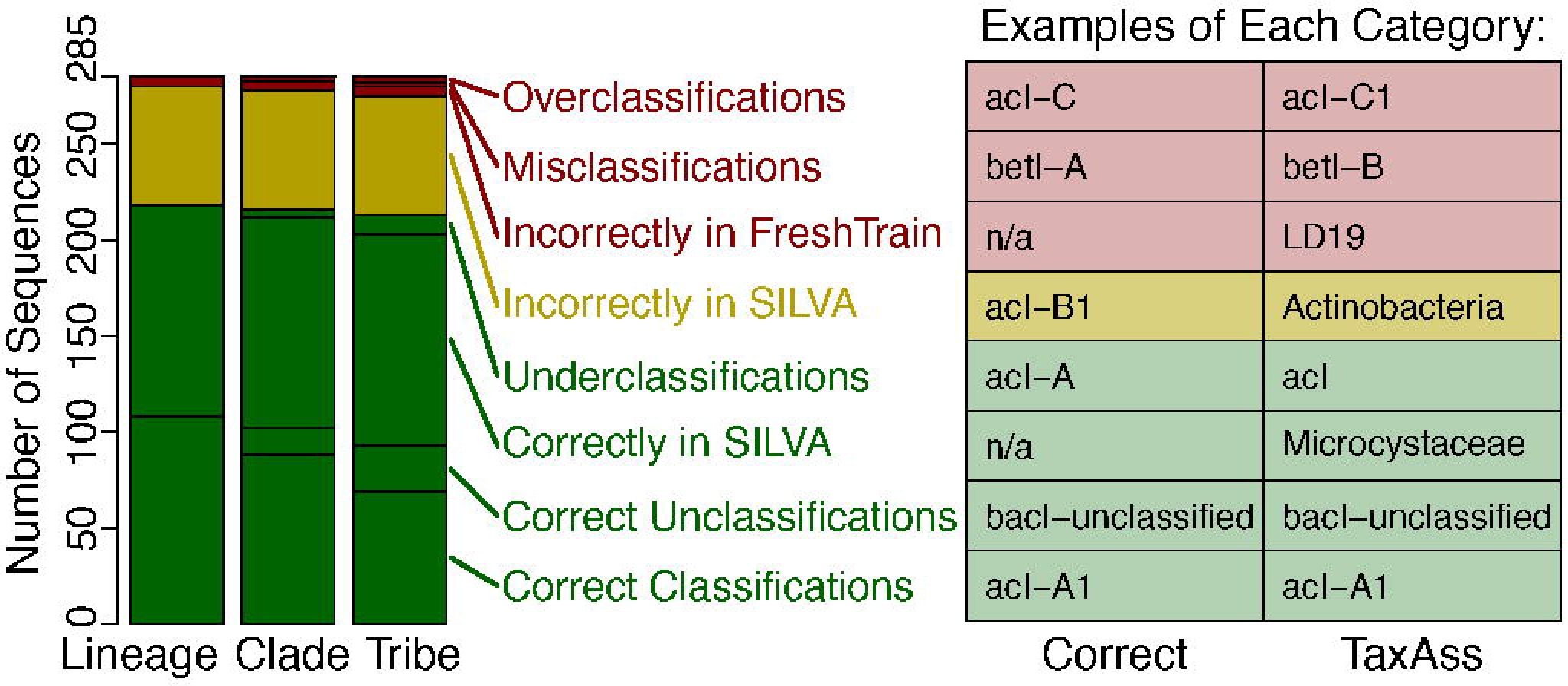
TaxAss validation with tags simulated from full-length Marathonas Reservoir clone libraries. Tags simulated by trimming full-length sequences to the V4 region were classified by TaxAss, and the resulting classifications were compared to “correct” classifications determined by manually aligning the full-length sequences to the FreshTrain. **A and B:** Correct classifications are in green, lost ecosystem-specific classifications are in yellow, and incorrect classifications are in red. **A:** Number of unique sequences in each classification category at fine-resolution taxa levels. **B:** Examples of classifications that fit into each classification category. Tabular results from this and additional amplicon region simulations are available in Supplemental Table 1.

We found that the majority (74.7 %) of V4 tag sequences were classified correctly at the species/tribe taxa level and that 86 % of the incorrect assignments were due to sequences being classified using Silva when they should have received FreshTrain nomenclature, which results in correct, though not ecosystem-specific, classifications. The remaining incorrect assignments stemmed from overclassification errors (1.1 %), misclassification errors (0.7 %), or incorrect inclusion in the FreshTrain classification set (1.8 %), which can result in overclassification, misclassification, or underclassification. (Figure 2a). Examples of each classification category are shown in the table in Figure 2b. We do not consider underclassifications to be an error because underclassifications are expected due to the lower phylogenetic resolution of short tag sequences compared to full-length sequences. We found slightly lower error rates for the longer V4-V5 and V3-V4 amplicon regions (Supplemental Table 1).

### Fine-Resolution Classifications Increased

To test whether TaxAss improved taxonomic classification over solely using a comprehensive database, we assigned taxonomy to a Lake Mendota amplicon dataset first by using Silva alone and then by using TaxAss to leverage both Silva and the FreshTrain (Figure 3a and Table 1). We compared the percent of reads classified by both methods and observed a marked improvement in the percent of the dataset classified to the fine taxa levels of family/lineage, genus/clade, and species/tribe. At species/tribe level, the percent of reads classified increased from 0% to 41%, at genus/clade level they increased from 35% to 63%, and at family/lineage level they increased from 72% to 82%. In addition to these increases in classifications, TaxAss also improved the quality of classifications because the FreshTrain is curated with terminology and phylogeny consistent with the freshwater microbial ecology literature. For example, the abundant and cosmopolitan freshwater tribe acI-A1 is split into hgcl clade and *Candidatus* Planktophila in Silva; acI-A4 and -A5 are also grouped with Silva’s hgcl clade, and acI-A3 is grouped with *C.* Planktophila. At family/lineage level, Silva alone could classify a majority of the dataset, but 72% of those Silva-classified reads received more meaningful ecosystem-specific nomenclature when using TaxAss.

**Figure 3.**
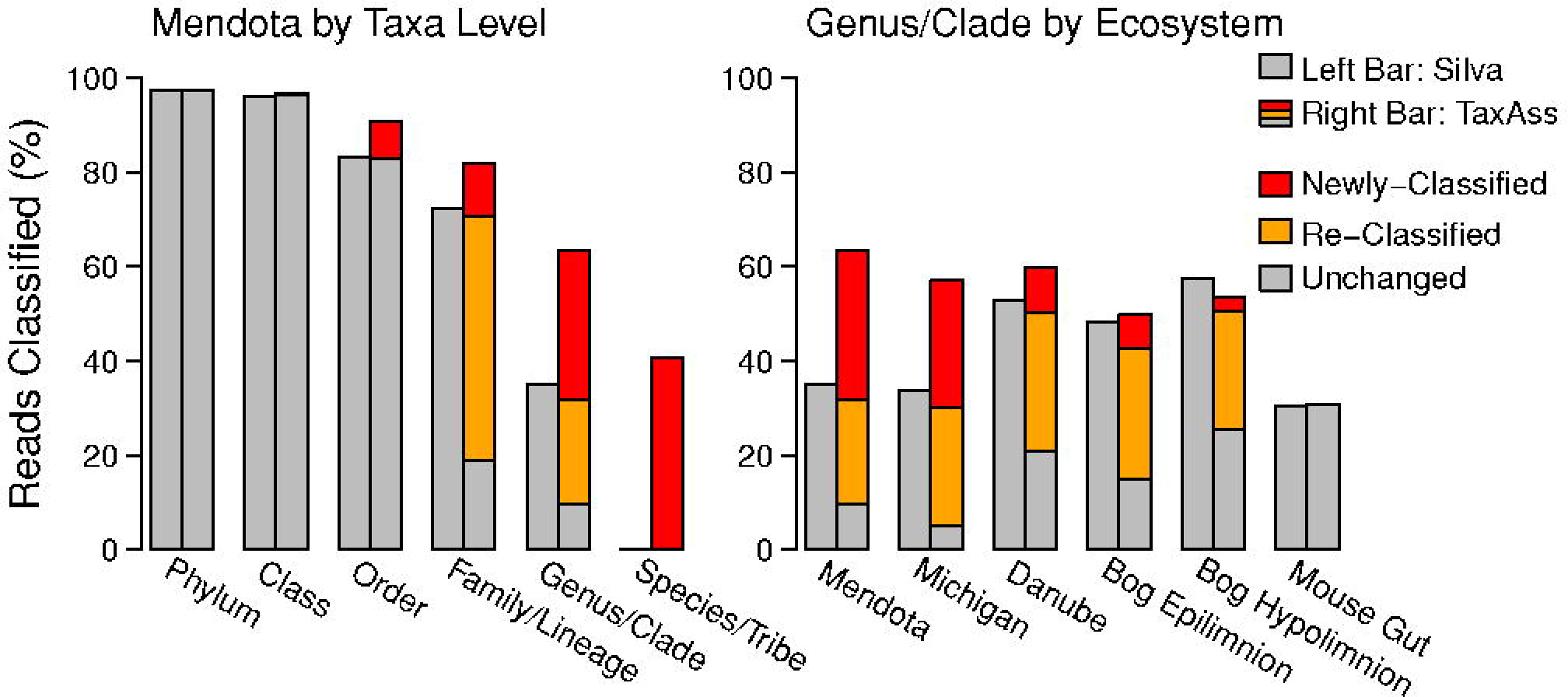
TaxAss performance compared to Silva-only performance. **A and B:** Left bars represent the Silva-only classification and right bars represent the TaxAss classification that leveraged both Silva and the FreshTrain. Within the right bars, red reads were classified by the FreshTrain using TaxAss and were left unclassified using only Silva; yellow reads were classified by the FreshTrain using TaxAss but received Silva classifications using only Silva, and grey reads were classified by Silva when using TaxAss.
**A:** In the Lake Mendota dataset, TaxAss leveraged the FreshTrain and Silva to achieve improved fine-resolution classifications. **B:** TaxAss achieved improvements in a range of freshwater datasets despite the FreshTrain’s primary focus on temperate lake epilimnia. Few changes in classification were observed in the mouse gut control. Versions of this figure across all datasets and taxa levels can be found in Supplemental Figure 1.

**Table 1.**
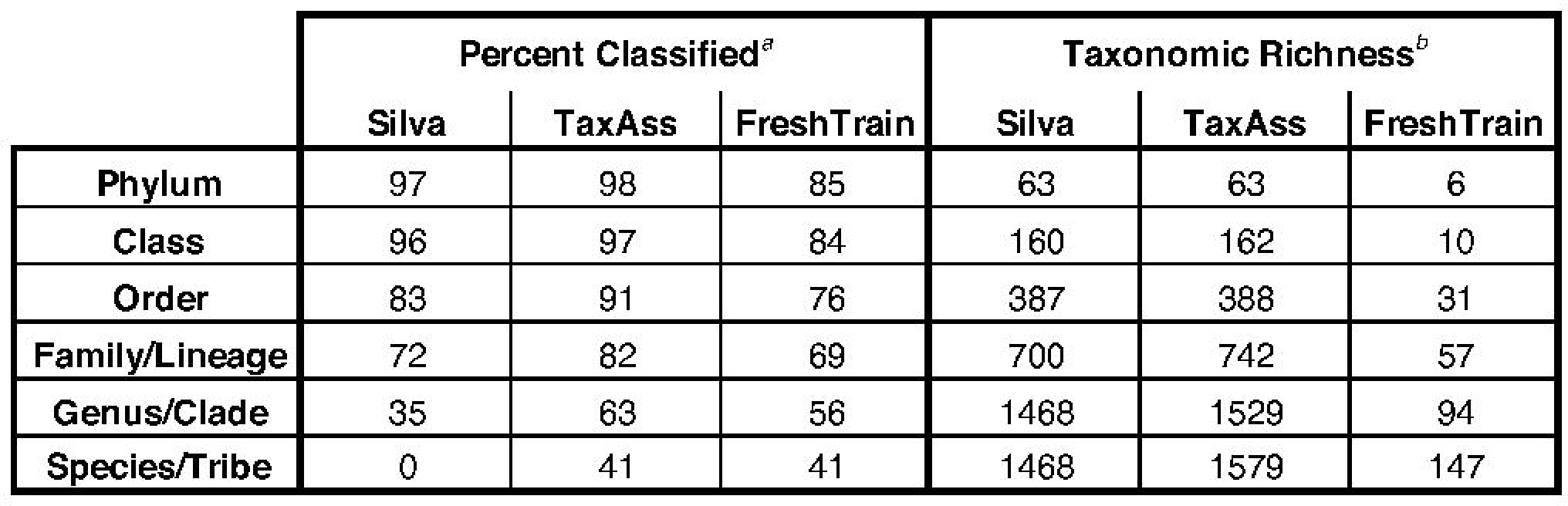
Classification of the Lake Mendota dataset* using Silva alone, the FreshTrain alone, or using TaxAss to leverage both databases. ^*a*^Percent classified = total reads classified / total reads in dataset x 100% ^*b*^Taxonomic Richness = total unique classifications *Versions of this table for each tested dataset can be found as Supplemental Table 2.

|  | Percent Classified<sup>a</sup> |  |  | Taxonomic Richness<sup>b</sup> |  |  |
| --- | --- | --- | --- | --- | --- | --- |
|  | Silva | TaxAss | FreshTrain | Silva | TaxAss | FreshTrain |
| **Phylum** | 97 | 98 | 85 | 63 | 63 | 6 |
| **Class** | 96 | 97 | 84 | 160 | 162 | 10 |
| **Order** | 83 | 91 | 76 | 387 | 388 | 31 |
| **Family/Lineage** | 72 | 82 | 69 | 700 | 742 | 57 |
| **Genus/Clade** | 35 | 63 | 56 | 1468 | 1529 | 94 |
| **Species/Tribe** | 0 | 41 | 41 | 1468 | 1579 | 147 |

The FreshTrain reference sequences come exclusively from temperate lake epilimnia, and many of them come from Lake Mendota itself. Lake Mendota is a eutrophic, temperate lake in Wisconsin, USA, and the Lake Mendota amplicon dataset consists of 95 epilimnetic samples collected by the North Temperate Lakes Microbial Observatory over 11 years. To test TaxAss’s efficacy when the ecosystem-specific database is less representative of the ecosystem under investigation, we classified amplicon datasets from a range of freshwater ecosystems first by using Silva alone and then by using TaxAss to leverage Silva and the FreshTrain (Figure 3b). The additional ecosystems we chose included the epilimnion of oligotrophic Lake Michigan (20), the eutrophic Danube River (21), and the epilimnion and hypolimnion of dystrophic Trout Bog (WI, USA) (22). We also used a mouse gut dataset (23) as a negative control to ensure that TaxAss would not assign FreshTrain classifications erroneously. All freshwater datasets showed improvements at all fine taxa levels (Figure 3b and Supplemental Figure 1), with the amount of improvement reflecting the similarity of each ecosystem to the FreshTrain reference sequences. For example, the temperate Lake Mendota and Lake Michigan epilimnia received the most FreshTrain classifications (54 and 52% of total reads at genus/clade level), while the dystrophic bog hypolimnion benefited least (28% at genus/clade level). Only 0.1% of the mouse gut control dataset received FreshTrain classifications at the species, genus, or family levels.

### Richness Maintained

To test whether TaxAss improved taxonomic classification over solely using an ecosystem-specific database, we assigned taxonomy to the Lake Mendota dataset first by using the FreshTrain alone and then by using TaxAss to leverage both the FreshTrain and Silva. TaxAss maintained taxonomic richness at all taxa levels by classifying OTUs into a larger variety of taxonomic names (Figure 4 and Table 1). At the same time TaxAss prevented overclassifications and misclassifications at fine-resolution taxa levels compared to FreshTrain-only classifications (Figure 4b).

**Figure 4.**
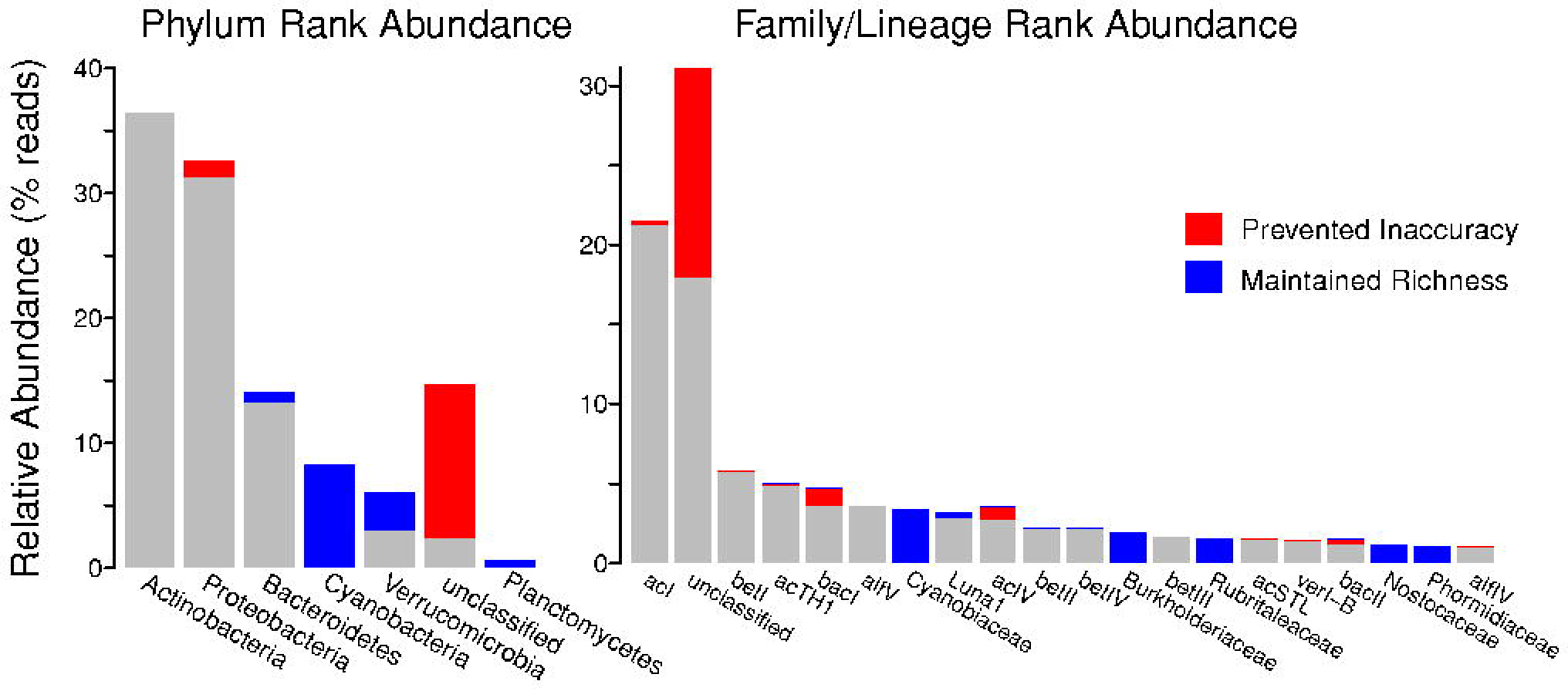
TaxAss performance compared to FreshTrain-only performance. **A and B:** Lake Mendota reads represented by blue bars were incorrectly classified as red bars in the FreshTrain-only classification. Rank order of the bars follows the TaxAss-classification rank abundances. Only taxa with at least 0.5% relative abundance are included, and at lineage level the number of bars displayed is further truncated to 20. **A:** TaxAss maintained phylum richness (blue bars) by classifying phyla using Silva when they are not included in the FreshTrain. **B**: TaxAss prevented lineage-level inaccuracies from misclassifications and overclassifications (red bars over known taxa), and lineage-level underclassifications (red bars over “unclassified” taxa). Versions of this figure across all test-ecosystems can be found in Supplemental Figure 2.

The FreshTrain is a more specific database with less taxonomic richness than Silva, so a decrease in taxonomic richness in a FreshTrain-only classification was expected. For example, the FreshTrain focuses on heterotrophic bacteria and does not include any Cyanobacteria, which comprised 8.3% of the Lake Mendota dataset. All of Lake Mendota’s cyanobacterial OTUs were classified as something else (99.9% as unclassified), which resulted in a loss of phylum-level richness in the FreshTrain-only classification (Figure 4a). In contrast, TaxAss maintained the taxonomic richness of a Silva-only classification (Table 1, Supplemental Table 2).

We also observed that some OTUs that TaxAss classified using Silva were misclassified or overclassified by the FreshTrain-only approach (Figure 4b). These incorrect classifications by the small FreshTrain database were less common than underclassification errors, but had significant effects on taxa relative abundances at finer-resolution taxa levels. Lake Mendota’s 5th most abundant lineage, the *Bacteroidetes* bacI, gained 30% more reads in a FreshTrain-only classification compared to using TaxAss. The classification errors TaxAss prevented were significant enough to change basic attributes such as rank abundances of top taxa, and had an even larger impact on the freshwater test-datasets that differed more from the FreshTrain references (Supplemental Figure 2).

### Percent Identity Cutoff

An OTU is classified taxonomically in the ecosystem-specific database only if it matches a sequence in that database at a sequence identity above the threshold set by the user. Therefore, the percent identity cutoff choice for taxonomic classification is central to the proper functioning of TaxAss because it determines which OTUs are classified in each database (ecosystem-specific vs comprehensive). If the percent identity cutoff is set too high, ecosystem-specific OTUs are passed to the comprehensive database for classification; while if it is set too low, non-ecosystem-specific OTUs are passed to the ecosystem-specific database for classification. In both scenarios the majority of misplaced OTUs will be unclassified at fine taxa levels. TaxAss allows users to compare the percent of reads classified with different percent identity cutoffs, the idea being that a percent identity that maximizes reads classified has minimized misplacement errors of the abundant OTUs (Figure 5).

We found that a percent identity cutoff of 98-99% was appropriate for the analyzed freshwater datasets, and we applied a cutoff of 98% when processing all data used in this paper (Figure 5). TaxAss allows users to choose a cutoff specific to their data by generating the plots shown in Figure 5, but users who wish to save computational time can simply choose a percent identity cutoff and only run the classification once.

**Figure 5.**
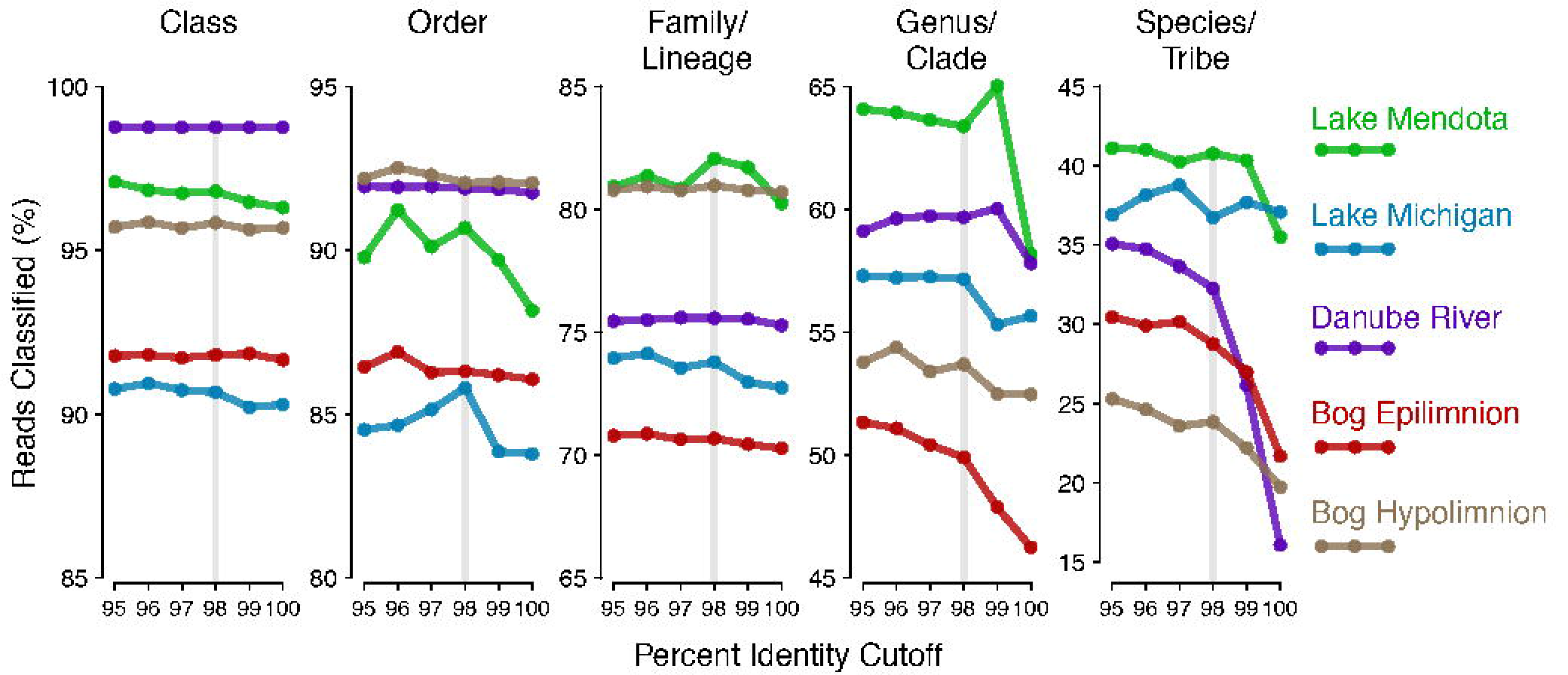
Percent identity where classifications are maximized. The percent of reads classified when using different percent identity cutoffs to separate out ecosystem-specific OTUs, shown for each freshwater dataset across taxa levels. Faint vertical lines highlight the 98 percent identity chosen for the analyses in this paper. OTUs are predominantly unclassified at fine resolution if they are placed in the wrong classification group, so this visualization is generated by TaxAss to help users choose a percent identity cutoff appropriate for their dataset.

### BLAST Conversion

The calculation of percent identity for use in database selection is based on the percent identity returned by The National Center for Biotechnology Information’s Basic Local Alignment Search Tool (BLAST) (24). The default megablast settings are appropriate for our application because they have been highly optimized to find short, highly similar alignments. However, BLAST finds areas of local similarity and there is no way to require BLAST to align the entire length of a query OTU’s sequence. 16S rRNA gene amplicon sequences are highly similar, and differences in taxonomic classification can be based on even a single mismatch in the amplified region. Therefore, we recalculated the percent identities BLAST returned into “full-length” percent identities for the entire query OTU’s sequence (Supplemental Document 1 and Equation 1).

We found that recalculating percent identity was necessary to prevent dissimilar OTUs from inclusion in the ecosystem-specific classification. For example, the FreshTrain does not include any reference sequences from the major freshwater phylum *Cyanobacteria*, so no cyanobacterial OTUs have high true percent identities to any references in the FreshTrain. We found that the percent identity recalculation was necessary to prevent some cyanobacterial OTUs from meeting the percent identity cutoff due to the original BLAST percent identities being based on only a short aligned section of the OTU sequence (Supplemental Figure 3).

We also found that it was necessary to recalculate the percent identity from several BLAST alignments (“hits”) for each OTU because the best BLAST hit did not always have the highest recalculated percent identity. TaxAss examines the top five BLAST hits, recalculates the percent identity of each, and then uses the highest recalculated percent identity to determine if an OTU meets the cutoff. To ensure enough BLAST hits were examined to consistently arrive at the highest possible recalculated percent identity, we calculated the proportion of times each BLAST hit number had the highest recalculated percent identity. In the Lake Mendota amplicon dataset the first BLAST hit almost always also had the best recalculated score, and the contribution of additional BLAST hits was very low, especially when only “good” hits above a stringent percent identity cutoff were considered (Table 2). In the Lake Mendota dataset at the chosen 98 percent identity cutoff, 99.68% of the best hits found by BLAST were also the best recalculated hits and only 0.07% of BLAST’s 5th hits were used. TaxAss generates a version of Table 2 for users’ individual datasets, and if they observe more high-number BLAST hits contributing to the best re-calculated hit they can increase the number of BLAST results used for the calculation.

**Table 2.** Agreement between BLAST and recalculated percent identities. ^*a*^Calculations performed only on sequences above listed recalculated percent identities ^*b*^*e.g.* 97.8% of BLAST’s first hits also had the highest recalculated percent identity

| BLAST<br>result | Percent Identity Cutoff Applied<sup>2</sup> |  |  |  |  |  |
| --- | --- | --- | --- | --- | --- | --- |
|  | 95 | 96 | 97 | 98 | 99 | 100 |
| Hit 1 | 96.8<sup>b</sup> | 97.8 | 99.0 | 99.68 | 99.90 | 100 |
| Hit 2 | 1.4 | 1.0 | 0.5 | 0.18 | 0.07 | 0 |
| Hit 3 | 0.5 | 0.3 | 0.1 | 0.04 | 0.01 | 0 |
| Hit 4 | 0.7 | 0.4 | 0.1 | 0.03 | 0.01 | 0 |
| Hit 5 | 0.7 | 0.5 | 0.2 | 0.07 | 0.01 | 0 |

## DISCUSSION

### Ecosystem-Specific Databases

The need for curated ecosystem-specific databases has been recognized by microbial ecologists studying many ecosystems. TaxAss was developed specifically to leverage the Freshwater Training Set (FreshTrain) (15), but it could be applied to custom databases curated for other ecosystems: the dictyopteran gut microbiota reference database (DictDb) (13), the rumen and intestinal methanogen database (RIM-DB) (11), the honey bee database (HBDB) (12), the microbial database for activated sludge (MiDAS) (14), and the human oral microbiome database (HOMD) (10). These databases were created by starting with a comprehensive database such as SILVA or Greengenes and then re-curating the reference sequences from the study ecosystem, sometimes also incorporating new reference sequences. Often during curation, phylogenies were collapsed to be monophyletic and incorporate new organisms, and abundant but unnamed organisms were given placeholder names to allow for consistent terminology among researchers.

DictDB, HBDB, and MiDAS are fully integrated with modified versions of the entire SILVA database, so a workflow like TaxAss that leverages two databases is not needed because the single merged database can be used in one step for taxonomy assignment. However fully integrated databases can be difficult to maintain over time because new versions of each database will diverge from each other, and TaxAss provides a means to circumvent this divergence. The FreshTrain is an example of this divergence in action. The FreshTrain was originally integrated into the Hugenholtz database that eventually became Greengenes, and Greengenes was last updated in May 2013. In addition, SILVA now contains more total references and has been updated as recently as December 2017, so some researchers prefer to use the more recently updated SILVA as their comprehensive database. Similarly, the FreshTrain has been updated almost annually since its creation as new full-length 16S rRNA gene sequences from freshwater ecosystems became available. TaxAss allows microbial ecologists to use the most up-to-date versions of their preferred databases without performing or waiting for reconciliation of each release.

Once an ecosystem-specific database has diverged from the comprehensive database, as occurred with the FreshTrain, leveraging the ecosystem-specific database for taxonomy assignment is no longer straightforward. Reintegrating the ecosystem-specific database into the comprehensive database is more involved than simply concatenating databases and removing duplicated references because conflicting phylogenetic structures must be resolved. Analysis of community amplicon data is a fairly routine part of many studies for which extensive phylogenetic curation would fall outside the scope. The FreshTrain has been used in a variety of ways since it diverged from the current version of Greengenes, and it is often difficult to discern the specifics from cursory sentences in a paper’s methods section. TaxAss provides a well-documented and rigorously tested workflow to leverage two conflicting databases without extensive curation.

### Current FreshTrain Usage

The simplest way the FreshTrain has been used to assign taxonomy to amplicon datasets is as part of a separate, complementary analysis. For example, in a study of the River Thames Basin (25), FreshTrain and Greengenes classifications were displayed side by side and separate metrics such as diversity indices were calculated for each. However, the bulk of the taxonomic analyses were carried out at the coarse phylum level despite most abundant OTUs having FreshTrain nomenclature. When the FreshTrain is used independently, the loss of richness in taxonomic classifications (Figure 4) makes it difficult to use ecosystem-specific classifications for entire-dataset analyses. TaxAss provides ecosystem-specific classifications without loss of taxonomic richness, thus allowing for a single comprehensive analysis.

Another straightforward approach has been to classify amplicon datasets sequentially, first using the FreshTrain and then re-classifying the unclassified sequences using a comprehensive database. For example, in a study of Lake Erken, Sweden (26), OTUs were first classified with the FreshTrain, and then unclassified OTUs were reclassified using SILVA. While this approach allows for a single analysis, the initial classification of all sequences with the small FreshTrain database can cause overclassification and misclassification errors (Figure 4b). TaxAss prevents this by splitting the OTUs into two groups prior to classification.

These classification errors when using the FreshTrain to classify all OTUs were observed in a study of cyanobacterial blooms in Yanga Lake, Australia (27), where the authors observed that cyanobacterial OTUs were forced into heterotrophic classifications. To prevent this, Greengenes was used for an initial classification, then only OTUs assigned to phyla included in the FreshTrain were reclassified and renamed with confidently assigned FreshTrain nomenclature. This approach prevented the misclassification of Yanga Lake’s abundant cyanobacterial OTUs, but it would not prevent overclassification of OTUs that belong to phyla included in the FreshTrain (Verrucomicrobia, Bacteroidetes, Proteobacteria, and Actinobacteria). In freshwater datasets such as bogs, rivers, and lake hypolimnia many organisms belonging to FreshTrain phyla differ significantly from the lake epilimnion references included in the FreshTrain. TaxAss prevents overclassifying and misclassifying OTUs of any phyla.

Another way to avoid the forcing observed with the Wang classifier is to use BLAST-based taxonomy assignment algorithms that determine assignments based on sequence similarity. Since the BLAST algorithm calculates an absolute similarity instead of a relative one, a similarity cutoff prevents classifications to dissimilar sequences. The BLAST method to assign taxonomy has been used with the FreshTrain to classify sequences from boreal lakes in Quebec, Canada (28). However, unlike the Wang classifier, BLAST only takes into account individual reference sequences and ignores their encompassing phylogenetic structure. The BLAST-based algorithm from Classification Resources for Environmental Sequence Tags (CREST) (29) addresses this by taking a lowest common ancestor approach. Each query OTU is classified to the finest taxa level that its top BLAST hits share. The CREST algorithm also has been used to assign taxonomy using the FreshTrain to sequences obtained from the Danube River in southeastern Europe (21). This approach avoided forcing and incorporated phylogenetic information in the taxonomy assignments, however it does not maintain diversity by also leveraging a comprehensive database. Additionally, the Wang classifier is more robust at coarser taxa levels and for shorter sequences (29), and it is implemented in common tools like mothur and QIIME. TaxAss allows users to leverage both ecosystem-specific and comprehensive databases using the highly trusted and conveniently implemented Wang classifier.

### Future TaxAss Usage

We recommend all microbial ecologists studying freshwater systems use the FreshTrain and TaxAss to classify their 16S rRNA gene amplicon datasets. This will result in a consistent, specific, and comparable vocabulary throughout the field, and will improve classification for analysis of individual datasets. We also recommend that microbial ecologists with different ecosystem-specific databases consider TaxAss when their databases diverge from the most up-to-date comprehensive database and phylogenetic curation is outside the scope of their project.

We recommend microbial ecologists create ecosystem-specific databases if one does not already exist, since they provide improved analysis and enhanced collaboration for the entire field. Phylogenies must be created from full-length 16S rRNA gene sequences, which are currently not collected as routinely as short amplicon sequences. However, we believe the benefit of these databases as a reference for the field and to improve taxonomic classification of amplicon sequences justifies the effort to create them, especially since TaxAss allows their use without constant re-curation. Additional full-length sequences to flesh-out the existing phylogenetic structure of organisms can be created with clone libraries, as was done for the FreshTrain. New sequencing technologies, such as the long reads produced by Nanopore (30) and PacBio (31, 32) instruments, promise even easier reference sequence generation in the future.

### Practical Guidance for Using TaxAss

TaxAss includes detailed descriptions of its constituent scripts including argument options and descriptions, so users are able to customize their analyses. The most important decision users make is the cutoff percent identity that determines which database is used to classify each OTU. If an OTU is above the cutoff (i.e. has high percent identity to an ecosystem-specific reference sequence) then it will be classified with the ecosystem-specific database. When the cutoff is higher, fewer OTUs are classified with the ecosystem-specific database and users run the risk of leaving some ecosystem-specific OTUs poorly classified by the comprehensive database. If the cutoff is lower, more OTUs will be classified with the ecosystem-specific database and users run the risk of overclassifications and misclassifications, and of losing taxonomic richness due to underclassifications. Users can decide on a percent identity cutoff at the beginning and run only one classification, or they can run TaxAss with several cutoffs and generate versions of Figure 5 to help guide their choice.

We found that a percent identity cutoff of 98-99% optimized classifications in our test-datasets. The finding that most OTUs match their ecosystem-specific reference sequences with such high percent identity suggests that the commonly chosen 97% sequence identity clustering is too coarse to observe fine taxa level dynamics. This is supported by previous findings that sequence identity-based OTUs can impose artificial delineations between organisms that affect results differently depending on the lineage (33), and that sequence identity-based OTUs can contain temporally discordant sequences (34). We recommend that users planning a taxonomy-centric analysis classify unique sequences after quality trimming and use fine-level taxonomic assignments to group their data instead of sequence-identity cutoffs. The classification step will take longer with a larger number of unique sequences, but users will likely save computational time overall by not clustering. For users who also want to emphasize traditional OTU-based analyses, we recommend choosing a finer sequence identity-based OTU definition such as 98 or 99% to best leverage the fine-level classification provided by TaxAss and a detailed ecosystem-specific database. When OTUs have been clustered based on sequence identity, we recommend that users choose the same or lower percent identity cutoff in TaxAss to prevent OTUs with constituent sequences falling on either side of the percent identity cutoff. We recommend a percent identity cutoff using similar metrics to those recommended for unique sequences when users define OTUs using other finely resolved techniques such as DADA2 denoising (5) or minimum entropy decomposition (3).

### TaxAss Informs Ecological Analyses

Taxonomy-based analyses allow researchers to compare results across datasets. Leveraging an ecosystem-specific database for taxonomy assignment results in a high proportion of fine resolution classifications, and grouping sequences based on these classifications is a dataset-independent way to describe community composition. The resulting taxonomic terminology is consistent and comparable between analyses, and the finely resolved taxonomic groupings enable ecologically informed analyses. TaxAss can complement OTU-based analyses independent of users’ chosen OTU definitions. Redefining OTUs to compare across datasets is computationally expensive, and is not possible for datasets created with differing amplification primers. When researchers use TaxAss to assign fine-level taxonomy to their datasets, colleagues can compare their results directly, without re-analysis and regardless of primer set. Additionally, taxonomic nomenclature can also bridge amplicon-based analyses and genomic analyses.

Leveraging ecosystem-specific databases for taxonomy assignment also improves researchers’ interpretations of individual datasets. Ecosystem-specific terminology is more meaningful because ecosystem-specific databases incorporate additional reference sequences, finer phylogenetic delineations, consistent nomenclature for uncultured organisms, and monophyletic structures. For example, the dominant lineage in freshwater is the FreshTrain’s actinobacterial lineage acI, which in Silva is usually classified as family Sporichthyaceae. Although a classification exists for this organism in both databases, the Silva family is much broader and also includes the separate FreshTrain lineages acSTL and acTH1. The FreshTrain’s finer-level phylogenetic information on these abundant freshwater actinobacteria is based on manually curated alignments and ecological information such as their occurrence in different lakes and is supported by prior work suggesting the clades and tribes are ecologically and metabolically differentiated (15, 35, 36). The fine-resolution taxonomy assignments provided by TaxAss and an ecosystem-specific database allow researchers to link their amplicon datasets with known ecophysiological traits.

Ecosystem-specific phylogenies that are not fully incorporated into a comprehensive database are not straightforward to leverage for taxonomy assignment. The FreshTrain, for example, has diverged from Greengenes since it was created, and it has been used for taxonomy assignment with inconsistent and sometimes unreliable methods. TaxAss is a well-documented, open source, and rigorously tested workflow that avoids the pitfalls of using a small database: forcing incorrect classifications onto sequences and losing taxonomic richness by leaving unrepresented organisms unclassified. At the same time, TaxAss achieves the benefits of an ecosystem-specific database: more meaningful nomenclature, larger proportions of the dataset classified, and finer-resolution classifications.

## METHODS

### How to Use TaxAss

TaxAss replaces only the taxonomy assignment step of users’ preferred amplicon dataset processing pipeline such as mothur or qiime. TaxAss consists of a series of scripts using R, Python, bash, mothur, and BLAST that are run from the terminal command line singly or as a batch file. The input to TaxAss is a quality controlled fasta file, and if users opt to run the optional percent identity cutoff metrics a relative abundance table is also required. The output of TaxAss is the fasta file’s sequence IDs followed by their 7-level taxonomy assignments. Scripts, step-by-step instructions, and detailed explanations of script argument options are available online at https://github.com/McMahonLab/TaxAss.

### Percent Identity Recalculation

The naïve Bayesian algorithm used for taxonomy assignment (the Wang classifier) (16) can overclassify or misclassify OTUs when a close match does not exist in a small reference database. TaxAss uses the well-accepted Wang classifier, but avoids classification errors resulting from the effects of a small database by only classifying sequences for which a close reference exists. The National Center for Biotechnology Information’s Basic Local Alignment Search Tool (BLAST) (37) is utilized to split the amplicon dataset into two groups prior to classification: one is classified with the ecosystem-specific database, the other with the comprehensive database.

Blastn queries each OTU sequence against the ecosystem-specific database using the default megablast settings, which are optimized to find highly similar matches between sequences longer than 30 bp (24). However, BLAST returns the percent identity of the highest scoring pair (the “pident”), which does not necessarily include the full length of the query OTU sequence. OTU sequences are highly similar; a single mismatch can change a classification, so mismatches at the ends of OTU sequences (in the “overhang”) that BLAST leaves out of the alignment must be included in the percent identity cutoff used for classification. Therefore, the BLAST pident is recalculated to a full-length percent identity with the following equation:

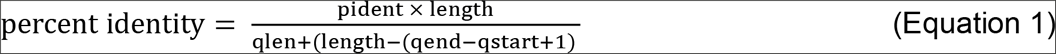

where “pident” is the percent identity returned by BLAST, “length” is the length of the alignment, “qlen” is the query length, “qend” is the query end, and qstart is the query start. All of these parameters are returned by BLAST output format 6, and detailed descriptions of what they are, the equation derivation, and an example alignment and calculation are included in Supplemental Document 1.

The recalculation to full-length percent identity is conservative; all query nucleotides not included in the alignment (nucleotides in the “overhang”) are considered mismatches. This means that it would be possible to exclude an OTU from the ecosystem-specific classification when its true percent identity is above the cutoff due to matches in unaligned overhangs. An example of this situation is illustrated in Supplemental Document 1. When the highest scoring BLAST alignments contain matches on the overhangs, some of the lower-scoring alignments will be longer, and therefore have a higher recalculated percent identity. To correct for this TaxAss recalculates the percent identity of the top five blast hits and uses the best one for the cutoff decision. TaxAss also shows users the distribution of chosen hits, so that settings can be re-evaluated if BLAST is not primarily returning hits that have the best recalculated percent identities.

### Cutoff Choice

OTUs with percent identities greater than or equal to the users’ specified cutoff are classified with the ecosystem-specific database using the Wang classifier as implemented by mothur. The remaining OTUs are classified with the comprehensive database, also using the Wang classifier. The choice of a percent identity cutoff is left to users so that they can balance their choices based on the structures of their datasets and their plans for analysis. If the percent identity cutoff is too low, dissimilar OTUs will be classified in the ecosystem-specific database and may be left unclassified or forced into incorrect classifications, but if the percent identity cutoff is too high, OTUs similar to the ecosystem-specific database will be classified by the comprehensive database and may end up poorly classified.

Users have the option to choose a cutoff percent identity at the start, or they can classify their datasets with multiple cutoffs and TaxAss will provide metrics to guide their decisions. These metrics include versions of Figure 5, which shows the cutoff choices that maximize the proportion of dataset classified at different taxa levels.

As an additional optional check, users can also classify their datasets with only the comprehensive database and then compare the classifications. TaxAss provides metrics to check for coarse-resolution misclassifications. Phylum- and class-level classifications are more reliable when assigned by a large comprehensive database that includes more diversity, so if ecosystem-specific classifications at these coarse taxa levels disagree with the comprehensive database’s assignments it suggests that the percent identity cutoff is too low. Only these coarse levels can be used to check for misclassifications because at finer taxonomic levels too many OTUs end up unclassified by the comprehensive database to compare assignments.

### Data Availability and Processing

The freshwater tag datasets used in this paper are all publicly available on the National Center for Biotechnology Information’s (NCBI) Sequence Read Archive (SRA). The accession numbers are: Lake Mendota (ERP016591) (34), Trout Bog (ERP016854) (22), Lake Michigan (SRP056973) (20), and Danube River (SRP045083) (21). The Lake Michigan and bog project accessions include additional sample types, so only the Lake Michigan and Trout Bog samples were used. The mouse gut dataset is the full version of the example data used by mothur’s miSeq SOP, and is available on the mothur website (https://www.mothur.org/wiki/MiSeq_SOP) (23). The Marathonas Reservoir clone library dataset is available from GenBank under accession numbers GQ340065−GQ340365 (19). The Marathonas Reservoir taxonomy determined by manual alignment to the FreshTrain is available from www.github.com/McMahonLab/TaxAss.

The taxonomy databases used in this paper are also publicly available. The Freshwater Training Set (FreshTrain) version used was FreshTrain30Apr2018SILVAv132 (15), which includes 1,318 freshwater heterotrophic bacterial references and is available from www.github.com/McMahonLab/TaxAss. The Silva database version used was version SSU 132 NR 99 (www.arb-silva.de) (8), which includes 213,119 bacterial and archaeal reference sequences clustered to 99 percent identity to avoid repeat references. A mothur-formatted version of this database obtained from www.mothur.org/wiki/Silva_reference_files was used for all analyses (accessed January 2018). Further details on download, versions, and formatting can be found in Supplemental Document 2 and in the detailed directions provided at www.github.com/McMahonLab/TaxAss.

Quality control of tag dataset fastq files was performed according to mothur’s MiSeq SOP (23, accessed September 2017) through the chimera checking step with mothur version 1.39.5 (17). The resulting unique sequences were defined as OTUs for all further analyses. The single-end sequencing datasets (Lake Mendota and Trout Bog) were also pre-processed with vsearch version 2.3.4_osx_x86_64 (38) to trim to uniform lengths and remove low quality sequences with > 0.5 expected errors. During TaxAss, the percent identity cutoff used for all datasets was 98%, and the Wang classifier’s bootstrap confidence was set at 80% for all classifications. Batch files that reproduce all download, quality control, and TaxAss processing for each dataset are available in Supplemental Document 2.

Manual alignment of the full-length Marathonas Reservoir clone library sequences to the FreshTrain database was performed using the program ARB (39). Chimeras were manually identified and removed from the analysis, and sequences without FreshTrain nomenclature were labelled unclassified. Tags were simulated by trimming full-length sequences to common primer regions with mothur version 1.39.5 (17). The primers used were V4 (515F: GTGCCAGCMGCCGCGGTAA, 806R: GGACTACHVGGGTWTCTAAT) (40), V4-V5 (515FB: GTGYCAGCMGCCGCGGTAA, 926R: CCGYCAATTYMTTTRAGTTT) (41), and V3-V4 (341F: CCTACGGGNGGCWGCAG, 805R: GACTACHVGGGTATCTAATCC) (42). A list of the processing commands used to trim sequences to the primer regions and classify them is available in Supplemental Document 3.

## ACKNOWLEDGEMENTS

This material is based upon work that is supported by the National Institute of Food and Agriculture, U.S. Department of Agriculture, under award number 2016-67012-24709 to JJH and WIS01789 to KDM. KDM also acknowledges funding from the United States National Science Foundation (NSF) Microbial Observatories program (MCB-0702395), the NSF Long Term Ecological Research program (NTL-LTER DEB-1440297), and an NSF INSPIRE award (DEB-1344254)

We thank the North Temperate Lakes Long Term Ecological Research Program (NTL-LTER) and the Microbial Observatory for their years of support collecting the Lake Mendota and Trout Bog time series. We also thank the Earth Microbiome Project for generously sequencing our Lake Mendota and Trout Bog datasets.

We thank Benjamin Crary and Jason Woodhouse for sharing insights from their efforts to combine taxonomy databases for classification, and the many users of TaxAss who have reported bugs during development. We thank Pat Schoss for his public review on bioRxiv and anonymous reviewer 2 for constructive and helpful feedback.

## LIST OF SUPPLEMENTAL DOCUMENTS

**Supplemental Table 1. Simulated V4, V4-V5, and V3-V4 tag classification accuracy as determined by comparison to their full-length alignment-based taxonomies.**

**Supplemental Table 2. Classification of all tested datasets using Silva alone, the FreshTrain alone, or using TaxAss to leverage both databases. (A version of Table 1 for each dataset used in this manuscript.)** ^*a*^Percent classified = total reads classified / total reads in dataset x 100% ^*b*^Taxonomic Richness = total unique classifications

**Supplemental Figure 1a. TaxAss compared to Silva-only performance. (A version of Figure 3a for each dataset used in this manuscript.)**

**Supplemental Figure 1b. TaxAss compared to Silva-only performance. (A version of Figure 3b for each taxa level.)**

**Supplemental Figure 2. TaxAss performance compared to FreshTrain-only performance. (A version of Figure 4 for each dataset used in this manuscript.)** Versions of Figure 4 for each test-dataset.

**Supplemental Figure 3. Importance of percent identity recalculation.** The phylum Cyanobacteria exemplifies why the percent identity recalculation is necessary. The ecosystem-specific FreshTrain database contains no cyanobacterial references, so cyanobacterial reads serve as a control for something that should be classified in the comprehensive Silva classification group. However, BLAST returned hits with high percent identities for cyanobacterial OTUs due to including short, high sequence identity partial alignments (red bars). After the TaxAss percent identity recalculation, the cyanobacterial OTUs had lower percent identities and none were included in the FreshTrain classification group (grey bars).

**Supplemental Document 1. BLAST percent identity recalculation.** This 3-page document defines the blast terminology, derives the equation to recalculate percent identity, and provides an example alignment and calculation.

**Supplemental Document 2. Data Processing Batch Files.** Directions for reproducing all data processing in this paper. These commands pair with a zip file hosted on the TaxAss github repo, which includes the folder structure and scripts used to download and process all tag datasets used in this manuscript‥ The folders include batch scripts that download each dataset, batch scripts that quality control each dataset, and batch scripts that perform TaxAss on each dataset, along with the versions of TaxAss scripts used in this paper.

**Supplemental Document 3. Tag Simulation Processing Commands.** This html file explains in more detail the TaxAss validation using tags simulated from full-length Marathonas clone library data. It also includes all commands necessary to reproduce the simulation.

